# Gastrointestinal Helminths Increase *Bordetella bronchiseptica* Shedding and Host Variation in Supershedding

**DOI:** 10.1101/2021.05.06.442912

**Authors:** Nhat Nguyen, Ashutosh K. Pathak, Isabella M. Cattadori

**Affiliations:** Center for Infectious Disease Dynamics, The Pennsylvania State University, University Park, PA 16802, USA; Department of Biology, The Pennsylvania State University, University Park, PA 16802, USA

## Abstract

Multi-species infections have been suggested to facilitate pathogen transmission and the emergence of supershedding events. However, how the interactions between co-infecting pathogens affect their dynamics of shedding, and how this is related to the host immune response, remains largely unclear. We used laboratory experiments and a modeling approach to examine temporal variations in the shedding of the respiratory bacterium *Bordetella bronchiseptica* in rabbits challenged with one or two gastrointestinal helminth species. Experimental data showed that rabbits co-infected with one or both helminths shed significantly more *B. bronchiseptica* by direct contact with an agar petri dish than rabbits with bacteria alone. There was also evidence of synergistic effects when both helminth species were present (triple infection). Co-infected hosts generated supershedding events of higher intensity and more frequently than hosts with no helminths. Model simulations revealed that the two helminths affected the relative contribution of neutrophils and specific IgA and IgG to *B. bronchiseptica* neutralization in the respiratory tract. In turn, these changes led to differences in the magnitude and duration of shedding among the various types of infection. However, the rapid variation in individual shedding, including supershedding, could not be explained by the interactions between infection and immune response at the scale of analysis that we used. We suggest that local rapid changes at the level of respiratory tissue probably played a more important role. This study provides novel insight into the role of helminths to the dynamics of respiratory infections and offers a quantitative explanation for the differences generated by two helminth species.

**Author summary:** The dynamics of bacterial infections can be facilitated by the presence of gastrointestinal helminths. Understanding the immunological processes that underline the pathogen-parasite interactions, and how they affect the dynamics of shedding, is important particularly for infections where control of the parasite maybe more effective than trying to reduce the bacterial infection. In this study, we examined the role of two gastrointestinal helminth species on the shedding of the respiratory *Bordetella bronchiseptica* using laboratory experiments of rabbits together with mathematical modeling. Hosts infected with helminths shed significantly more bacteria with evidence of supershedding, than hosts with only *B. bronchiseptica*. Simulations showed that by altering the relative contribution of neutrophils, specific IgA and IgG, helminths affected the control of bacterial infection in the respiratory tract. These interactions altered the intensity and duration of bacterial shedding, including the frequency and intensity of supershedding events. However, at the host level our model did not explain the rapid variation in shedding observed, suggesting that local processes in the respiratory tissue are critical for the prediction of the daily shed in the environment. This study advances our understanding of the dynamics of shedding in bacteria-helminth co-infections and provides insight that can be used to control disease spread.

## Introduction

Individual variation in pathogen transmission can increase the basic reproduction number *R*_0_ of a pathogen and determine whether an infection will invade and spread or stutter and quickly fades out in a population of susceptible hosts [1, 2]. One of the causes of this variation is associated with differences in the amount and duration of pathogen shedding, whereby some infected hosts shed disproportionately more and for longer than the average population, the so called supershedders [3, 4], while some others do not shed at all [5–7]. Co-infected hosts are frequently proposed to contribute to this variation, which could emerge from the immune-mediated interactions between pathogens and the consequences on host infectiousness [8–10].

While studies on the immunological response to multi-species infections has provided insight to the interactions between the host and its pathogens, there remains a need to identify how these processes relate to onward transmission, specifically the patterns of bacterial shedding. Dynamical mathematical models are particularly useful in disentangling these complexities as they can provide mechanistic-driven hypotheses that can be examined in relation to empirical data [11, 12]. In the current study, we applied this general approach to investigate the dynamics of shedding of the respiratory bacterium *Bordetella bronchiseptica* in rabbits experimentally co-infected with one or both of the gastrointestinal helminths *Trichostrongylus retortaeformis* and *Graphidium strigosum*. Specifically, we explored to what extent helminths could alter *B. bronchiseptica* shedding over time, whether the trend varied depending on the helminth species and to what extent the host immune response could explain the patterns observed.

*B. bronchiseptica* is a highly contagious bacterium of the respiratory tract that causes multiple symptoms and infects a wide range of mammals [13]. In rabbits and mice, and likely other mammal species, *B. bronchiseptica* is removed from the lower respiratory tract (lungs and trachea) via phagocytosis stimulated by a type 1 cell mediated antibody and neutrophil response [14–16]. Bacteria persist in the nose, although they are partially reduced by IgA antibodies in naïve and immunized mice [17, 18]. Transmission is poor among wild-type laboratory mice but increases among TLR4-deficient mice [19]. The TLR4-deficient response is associated with neutrophil infiltration, and the intensity of shedding has been found to be positively correlated with neutrophil counts. In contrast, rabbits naturally shed [20] and efficiently transmit *B. bronchiseptica* between animals [21]. The evidence of rapid outbreaks in pet kennels, livestock holdings and laboratory rabbittries are consistent with rapid transmission among animals in close contact. Occasionally, *B. bronchiseptica* spills-over into humans but there is no evidence of sustained onward transmission, as these human infections have invariably occurred in immuno-compromised individuals [13, 22]. Humans are primarily infected by the subspecies *B. pertussis* and *B. parapertussis* which are responsible for regular whooping cough outbreaks worldwide [23, 24]. Given the close relatedness between the subspecies, and considering the many similarities in the kinetics of infection and the immune response, *B. bronchiseptica* provides a good system to explore the dynamics of shedding in the Bordetella infections and the interaction with other infections, like gastrointestinal helminths.

In co-infections with other respiratory pathogens *B. bronchiseptica* contributes to exacerbate respiratory symptoms, including the development of acute pulmonary disease and bronchopneumonia, and ultimately host death [25–29]. Parasitic helminths commonly stimulate a type 2 immune response that interferes with the type 1 immune response developed against *B. bronchiseptica* and related subspecies [16, 30, 31]. In recent studies, we showed that rabbits infected with either *T. retortaeformis* or *G. strigosum* carried higher bacterial infections in the nasal cavity but there were no significant differences in the size and duration of infection in the lungs, when compared to rabbits infected with *B. bronchiseptica* alone [16, 20, 31]. These two helminths stimulate a similar type 2 response, however, while the former is reduced or cleared from the small intestine, the latter persists with high intensities in the stomach [32–34]. The modeling of the immune network in *B. bronchiseptica*-*T. retortaeformis* hosts and *B. bronchiseptica* infected rabbits highlights the important role of neutrophils and antibodies (IgA and IgG) to bacterial clearance from the lungs [16]. Most likely, this mechanism also explains the patterns observed in dual infections with *G. strigosum*, where bacteria are removed from the lower but not the upper respiratory tract [31].

To investigate the impact of gastrointestinal helminths to *B. bronchiseptica* shedding, we started by examining the pattern of shedding from four types of laboratory infections: i- *B. bronchiseptica* (B) only, ii- *B. bronchiseptica*-*G. strigosum* (BG), iii-*B. bronchiseptica*-*T. retortaeformis* (BT), and iv- *B. bronchiseptica*-*T. retortaeformis*-*G. strigosum* (BTG). We then developed an individual-based Bayesian framework that was applied to each of these four infections, independently. Specifically, we first built a dynamical model that investigated the relative contribution of neutrophils and specific IgA and IgG to bacterial infection in the whole respiratory tract. Second, we applied this dynamical model to laboratory data, where shedding was linked to the estimated bacterial infection. Helminths were assumed to alter the immune response to *B. bronchiseptica* by impacting the magnitude and time course of the three relevant immune variables. Simulations provided a mechanistic explanation of the kinetics of shedding and how the two helminth species, with contrasting dynamics, contributed to variation between and within types of infection.

## Results

### Experimental results: Helminth infections boost *B. bronchiseptica* shedding

For each type of infection, the number of bacteria shed, as determined by contact with an agar petri dish during fixed time (CFU/s), and the systemic immune response from blood samples, were collected every week or multiple times a week for every rabbit (See Material and Methods). There was large host variation in the level at which *B. bronchiseptica* was shed, both within and between the four types of infection (Fig 1). Some rabbits consistently shed large amounts of CFUs while others were low shedders or did not shed at all, specifically: 3 out of 16 for *B. bronchiseptica* only infected animals (B), 1 out of 23 for *B. bronchiseptica*-*G. strigosum* (BG), 3 out of 20 for *B. bronchiseptica*-*T. retortaeformis* (BT) and 0 out of 24 for the triple infection (BTG). These non-shedders were excluded from the subsequent analyses. High shedders were found during the first 60 days post infection although some of these continued to shed large numbers of bacteria well beyond this time (Fig 1 and 2). The level of shedding was significantly higher in rabbits co-infected with helminths than *B. bronchiseptica* only animals (pair-wise Wilcoxon test for all: *p* < 10^*−*4^). When we considered the co-infections, no significant differences were observed among the three groups although the triple infection did exhibit a tendency for the highest shedding intensity (BTG: 0.42, 95%CI: 0.29-0.52) followed by the BG group (median: 0.40, 95%CI: 0.30-0.50) and the BT rabbits (0.33, 95%CI: 0.23-0.49) (Fig 2).

**Fig 1.**
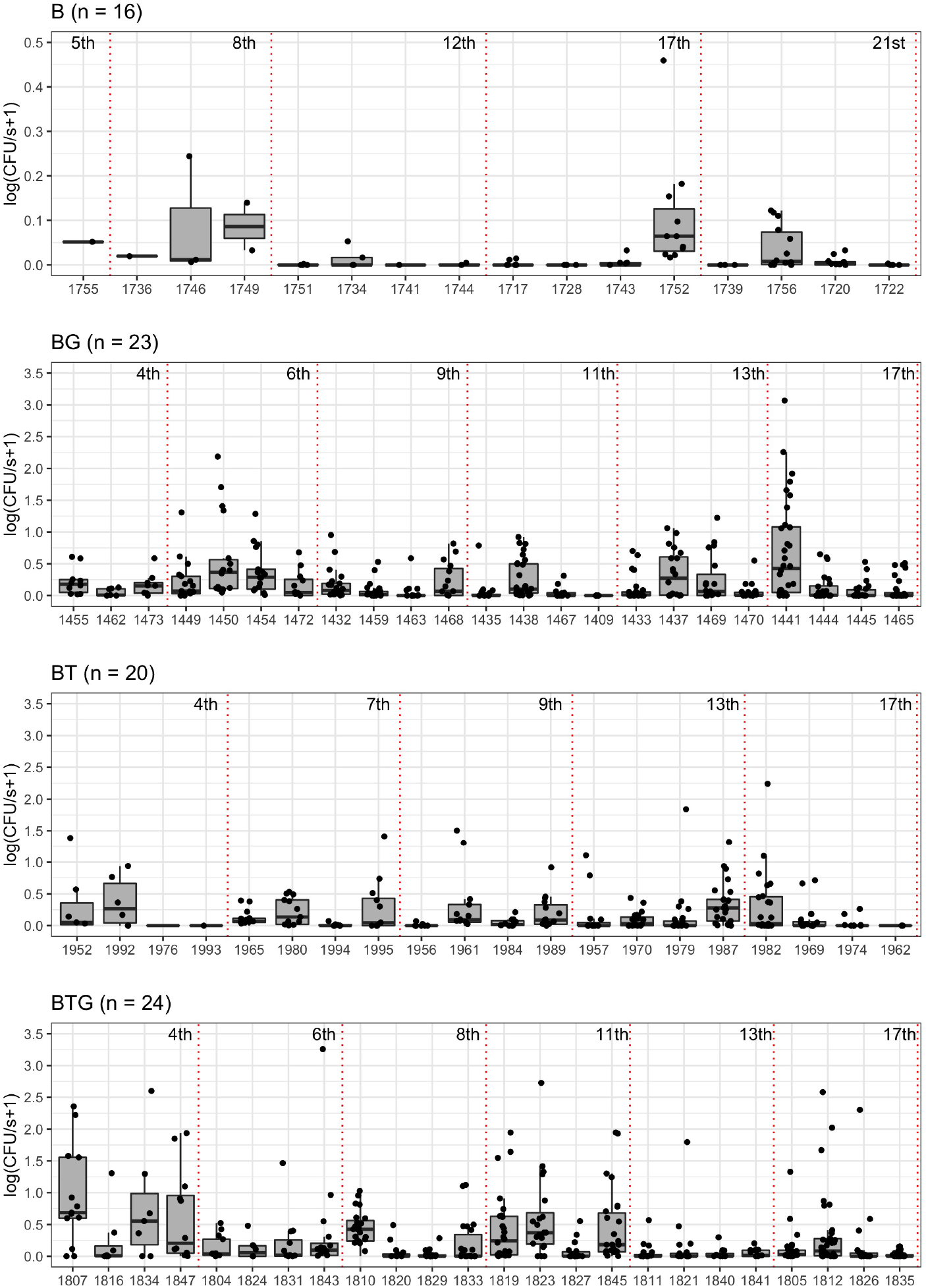
Observed intensity of *B. bronchiseptica* shed by contact with a Bordet-Gengou agar petri dish, and over a fixed time, by infected rabbit. The four types of infection are reported: *B. bronchiseptica* (B), *B. bronchiseptica*-*G. strigosum* (BG), *B. bronchiseptica*-*T. retortaeformis* (BT), and *B. bronchiseptica*-*T. retortaeformis*-*G. strigosum* (BTG) (top-down). For each infection, rabbits are ordered by their time of sacrifice (early/left, late/right) and experimental weeks are represented as dotted vertical lines. For every rabbit the following are reported: observed shedding events (black points), 25^*th*^ and 75^*th*^ percentiles (top and bottom hinges), median values (thick horizontal lines) and values within the 1.5 inter-quartile range (IQR, whiskers extendining up to 1.5*IQR). The total number of rabbits monitored in each infection is reported in parenthesis at the top left of each plot.

**Fig 2.**
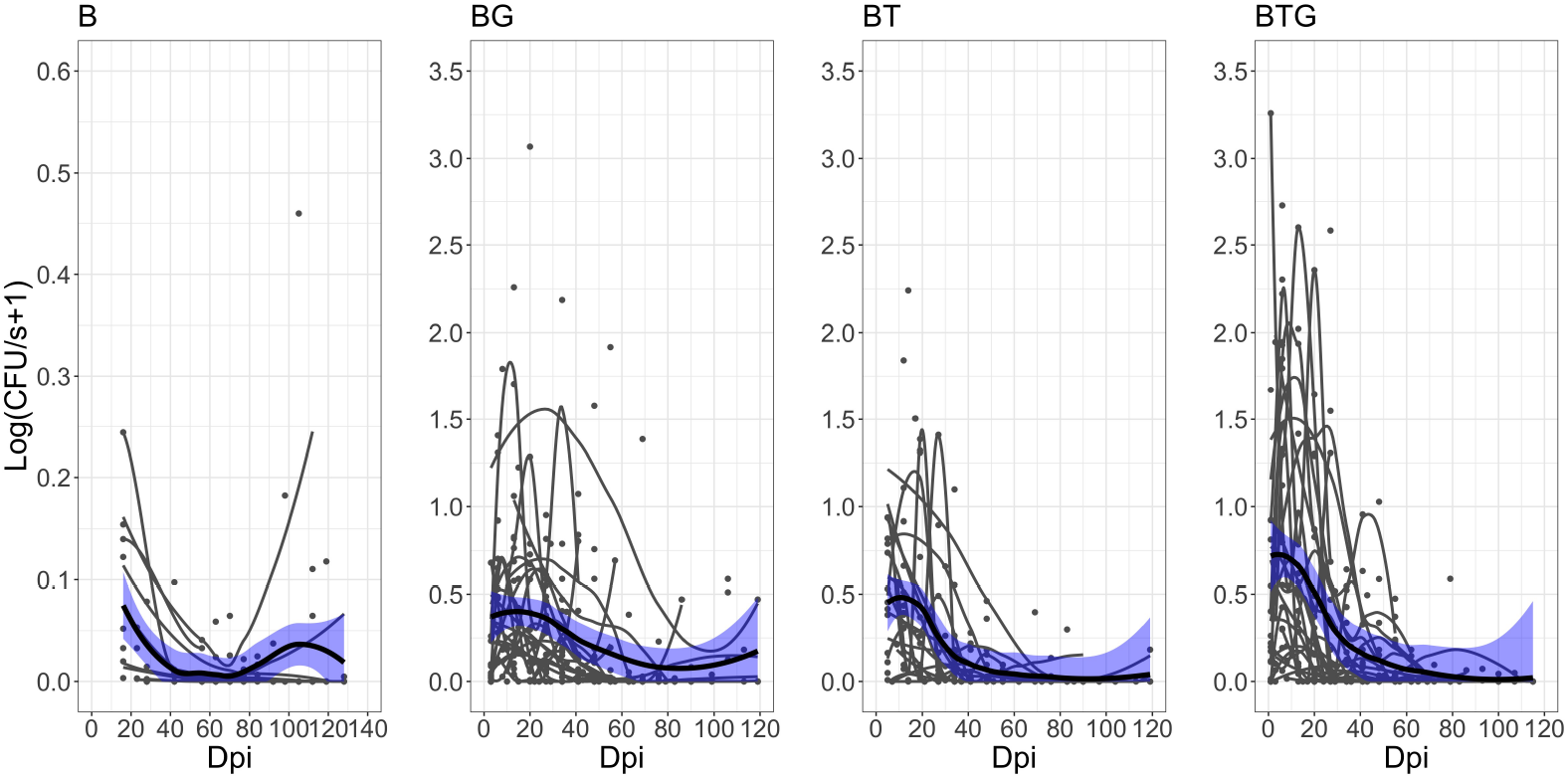
Observed *B. bronchiseptica* shed by Days Post Infection (DPI) for rabbits from the B, BG, BT and BTG group. Shedding events (black points), host individual trajectories (smoothed thin lines), and median population trend (smoothed thick lines) with 95% CI (blue areas) are reported. Host trajectories are reported only for time series with at least 4 shedding events. Note the much lower shedding values, and different y-axis, for the B panel. Further details in Fig 1.

Within groups many events were null, even though rabbits were known to be infected and interacted with a petri dish (Fig 3). Of these null events, at least 65% were recorded in the helminth-infected rabbits and increased to more than 80% in *B. bronchiseptica* only hosts (Fig 3). This supports the hypothesis that *B. bronchiseptica* is transmitted only during a small fraction of contacts, specifically, during 35%, or less, of the remaining shedding events, irrespective of the type of infection considered. Overall, BTG rabbits exhibited the largest variation in the intensity of shedding (log(CFU/s): mean=0.352, var=0.372) followed by the BG (mean=0.282, var=0.207) and the BT (mean=0.230, var=0.160) hosts, and last the *B. bronchiseptica* only infected rabbits (mean=0.028, var=0.004). Few events were characterized by high bursts, i.e. a large number of CFUs counted in a petri dish (hough rabbits were known to be infected and interacted with a petri dish (Fig 3 and 4). Therefore, while the percentage of shedding events by contact tend to be similar for single (about 20%) and helminth-infected rabbits (about 30%), the magnitude of these events is substantially higher for the latter group and should lead to a higher probability of onward transmission given the same number of contacts.

**Fig 3.**
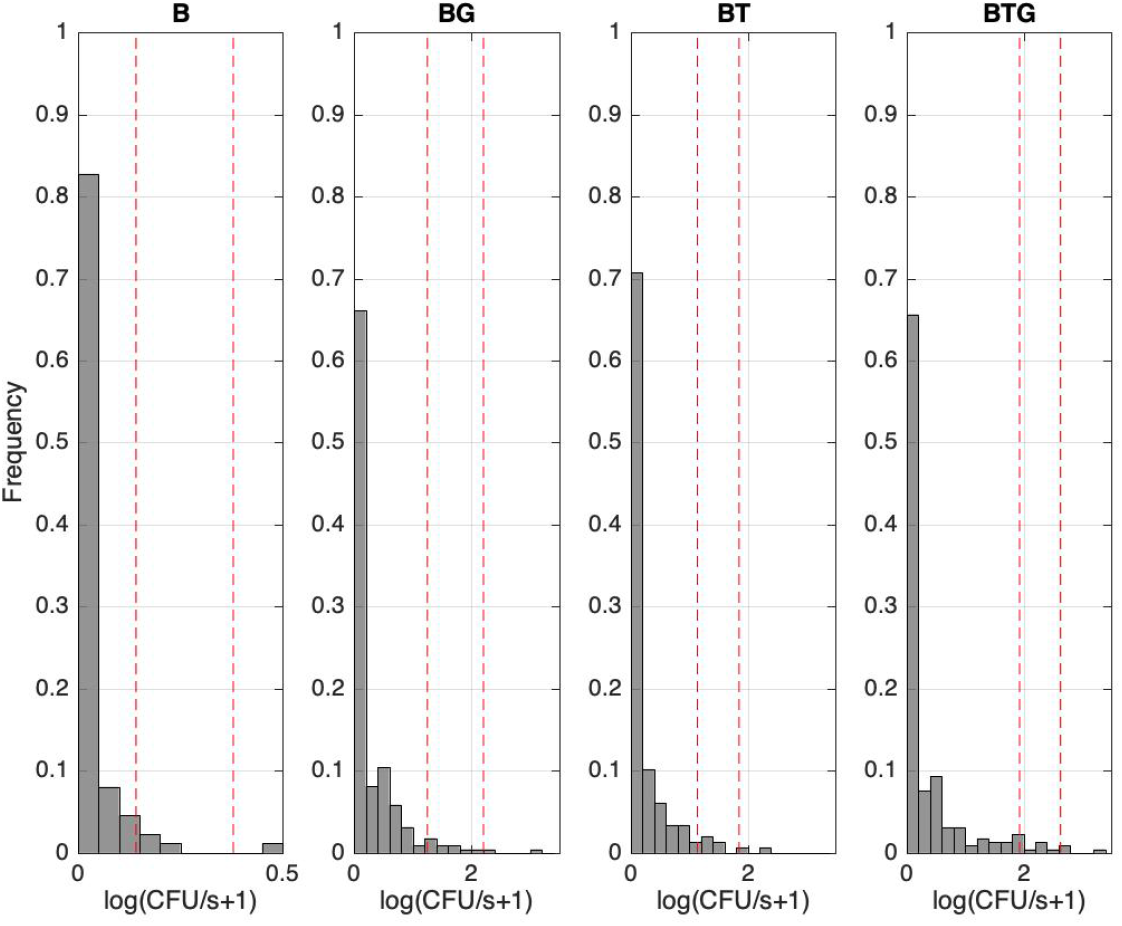
Frequency of observed *B. bronchiseptica* shedding by event intensity for B, BG, BT and BTG. Events have been groupped by shedding intensity using intervals of 0.2 log-unit for the co-infection groups and 0.05 for the B rabbits. Red lines indicate the 95^*th*^ and 99^*th*^ percentile thresholds, from left to right, respectively. Note the different x-axis, for the *B. bronchiseptica* only panel.

**Fig 4.**
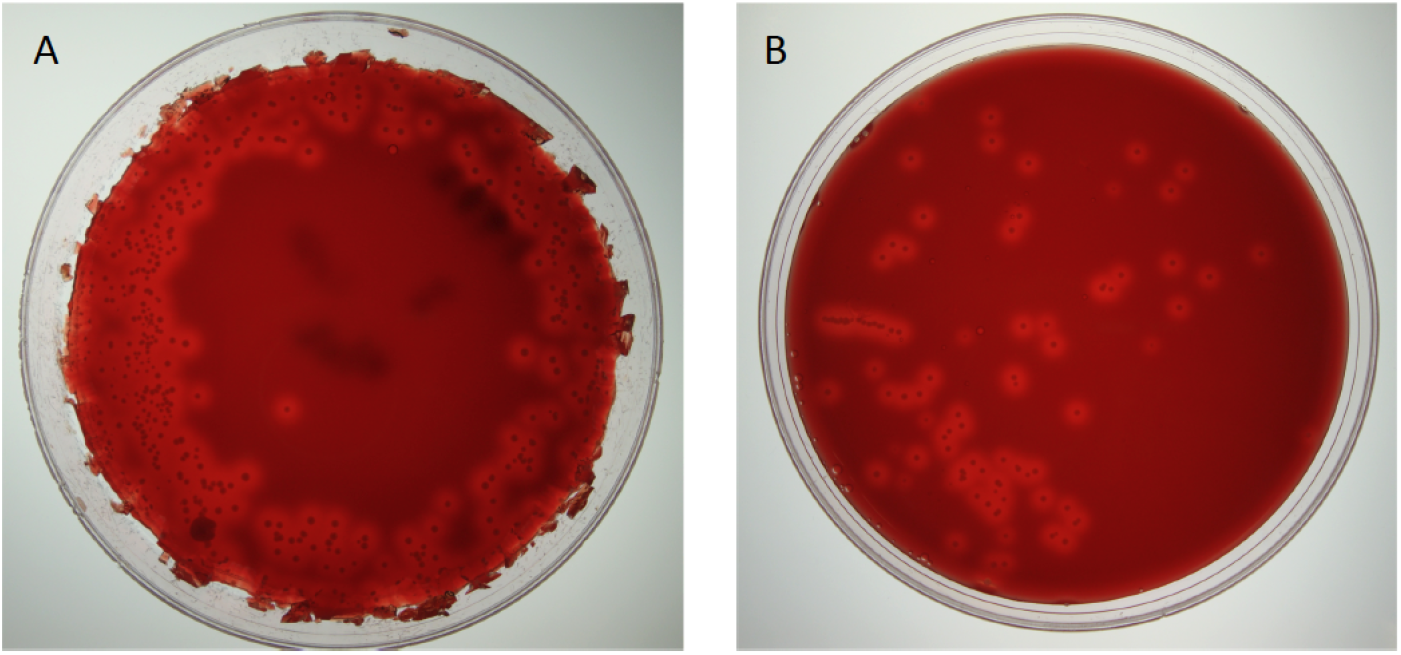
*B. bronchiseptica* shedding on BG-blood agar petri dishes. Examples of A: supershedding event and B: average shedding event.

### Experimental results: Helminth infections promote to *B. bronchiseptica* supershedding

The definition of supershedder, namely, the threshold above which there is evidence of a supershedding event, depends on the pathogen and the host, and it is usually based on the assumption that hosts carry one single infection [3]. If we estimate this threshold using the *B. bronchiseptica* only infected animals, and define supershedders as the hosts that have at least one shedding event above this threshold, for example the 99^*th*^ percentile, our cut-off value is log(CFU/s+1)=0.27 (Fig 3). This indicates that a limited 15% (2 out of 13) of the rabbits with *B. bronchiseptica* can be classified as supershedders. However, if we apply the same cut-off to the co-infected groups the percentage is significantly higher, specifically: 95% (21 out of 22) for the BG, 88% (15 out of 17) for the BT and 96% (23 out of 24) for the BTG group.

This threshold definition provides a common reference value for our types of infections, however, we show that it is dependent on host status (i.e. it is calculated using hosts with single infections), and is not representative in other settings, such that it overestimates the number of supershedders in hosts that are co-infected. A way to overcome some of the limitations of this approach is to estimate the cut-off value for each group of infected hosts independently. Therefore, for the co-infected rabbits, a 99^*th*^ percentile threshold applied to each infection provides the following values: log(CFU/s+1)=2.11 for BG, 1.66 for BT and 2.60 for BTG, and the fraction of hosts with, at least, one supershedding event above this value is now: 9% (2 out of 22) for BG, 12% (2 out of 17) for BT and 12% (3 out of 24) for BTG (Fig 3). This new estimation is based on the same general assumption for each group while considering the properties of the specific dataset. Similarly, if we select the less limiting 95^*th*^ percentile threshold, the number of rabbits with at least one supershedding event doubles, or increase even more, in some of the groups as: 23% (3 out of 13) for B, 23% (5 out of 22) for BG, 41% (7 out of 17) for BT and 37% (9 out of 24) for BTG (Fig 3). Likewise, the threshold value decreases to half for B and BG and less so for BT and BTG as: log(CFU/s+1)= 0.13 for B, 1.17 for BG, 1.10 for BT and 1.9 for BTG.

Consistent with the general pattern of shedding, supershedding events were found more often in the initial four/five weeks post infection (Fig 2), although later events were also observed, especially for the BG and BTG groups, and with some rabbits contributing multiple times. Importantly, since co-infected rabbits have a higher likelihood of becoming supershedders, there is an equally higher probability of onward transmission, given a contact, for these hosts compared to *B. bronchiseptica* only rabbits.

### Modeling results: Changes in the relative role of neutrophils and antibodies contribute to variation in *B. bronchiseptica* dynamics of infection between groups

We examined the hypothesis that *B. bronchiseptica* shedding is controlled by neutrophils, specific IgA and IgG produced against the bacterial infection in the respiratory tract, and helminths affect shedding by altering the magnitude and time course of the three immune variables. Our model formulation did not explicitly include the intensity of infection of the two helminths, rather, we examined how they altered the immune response to *B. bronchiseptica* and, with this, shedding. Here, we report on the relationships between immune responses and dynamics of infection by fitting a dynamical model to the experimental data, while in the next section we examine the relationship between predicted level of infection and estimated intensity of bacteria shedding. For the *B. bronchiseptica* only rabbits, model fitting was performed on the pooled data and did not include individual variation; this was because of the limited data on shedding available from these rabbits. The description of model framework and parameters calibration, including the sensitivity analysis of key parameters and posterior parameter estimates, are outlined in Material and Methods and Supplementary Information.

Model fitting captured the empirical trends of neutrophils and specific antibodies well, both as profiles representative of the whole group of hosts in each type of infection and also as profiles of every individual within each type of infection (Table 2, Fig 5 and S1). Consistently among the infections, there was rapid bacterial replication following the initial inoculum, the growth rate, *r*, was high for BTG (1.47, 95%CI [1.10–1.83]) and BG (1.18, 95%CI [0.83–1.54]), less so for BT (1.01, 95%CI [0.83–1.18]) and much lower in single infection (0.06, 95%CI [0.006–0.12]). *B. bronchiseptica* growth prompted a rapid immune response. In all four types of infection, neutrophils showed the fastest rate of increase, *a*, followed by specific IgA and then specific IgG (Table 2, Fig 5 and S1). Neutrophils were also the fastest to decrease, at a rate *b*, and to return to original values with the neutralization, albeit without complete clearance, of bacteria from the respiratory tract (Fig 5 and S1). As reported in our previous studies on this system, *B. bronchiseptica* is removed from lungs and trachea but persists in the nasal cavity [16, 20, 31]. The increase of specific IgA and IgG responses was slow, however, once IgG reached the asymptotic value at around 20 days post infection, it remained high throughout the trials, while IgA peaked around day 20 and decreased thereafter, although it had not fallen to baseline levels by the end of the trials (Fig 5 and S1).

**Table 1.**
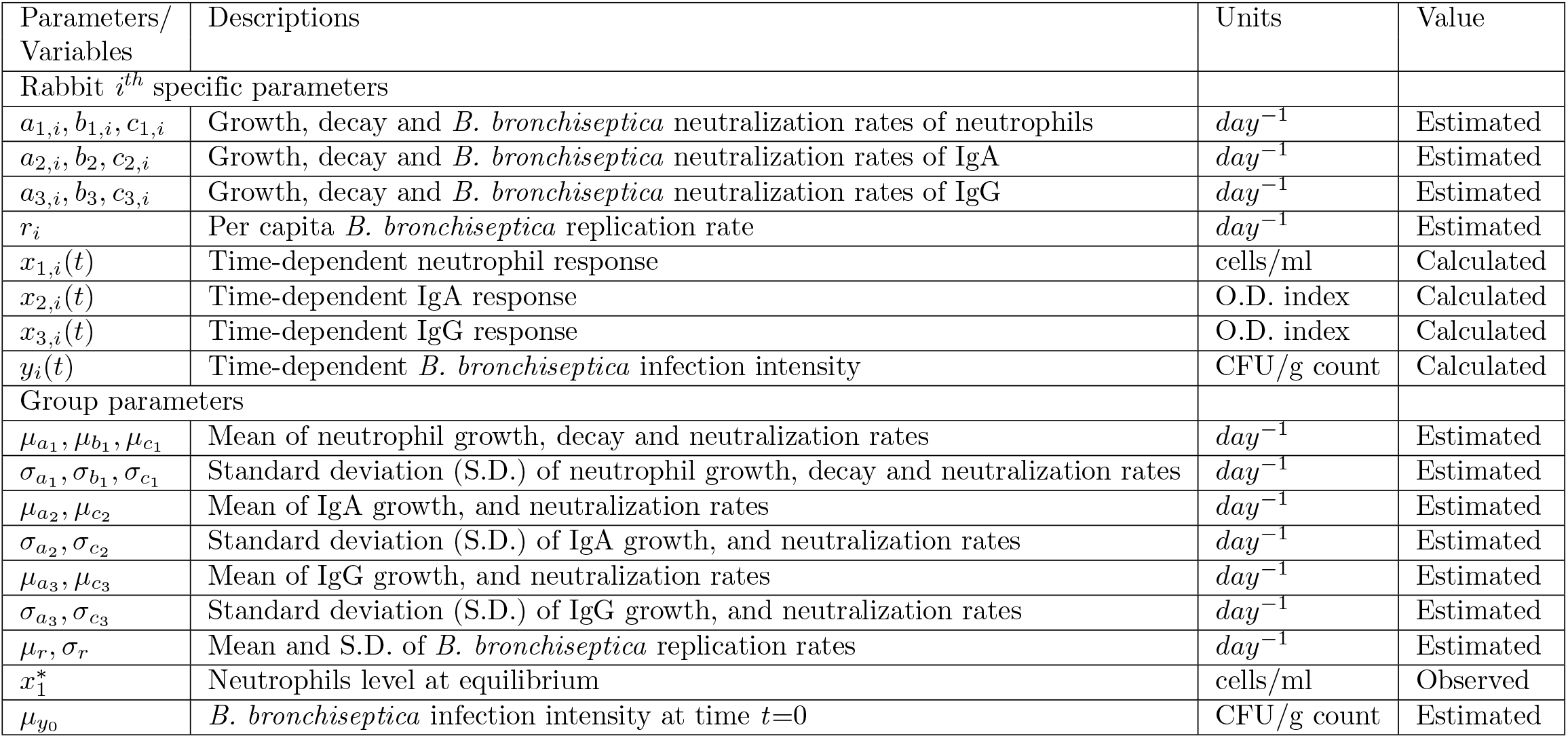
Description and unit of parameters and variables used in the dynamical and observed models.

**Table 2.** Estimated parameters of the host immune response and *B. bronchiseptica* infection at group level for the reduced model. Mean values, 95% CI and S.D. for each group are reported. The complete description of the parameters is provided in Table 1. For B group the parameters represent the fit from pooled data without individual variation.

| Parameters | B | BG | BT | BTG |
| --- | --- | --- | --- | --- |
| $\mu_{a_1} * 10^{-4}$ | 2.89 (1.17–4.61) | 1.08 (0.31–1.85) | 3.32 (1.73–4.90) | 0.93 (0.35–1.50) |
| $\mu_{a_2} * 10^{-5}$ | 2.0 (1.65–2.35) | 0.81 (0.51–1.11) | 0.99 (0.61–1.36) | 1.36 (0.91–1.81) |
| $\mu_{a_3} * 10^{-5}$ | 1.51 (1.27–1.75) | 0.66 (0.43–0.89) | 0.78 (0.49–1.36) | 0.86 (0.59–1.81) |
| $\mu_{b_1}$ | 1.20 (0.83–1.57) | 0.54 (0.37–0.71) | 0.86 (0.61–1.11) | 0.58 (0.40–0.77) |
| $b_2 * 10^{-3}$ | 7.07 (5.55–8.59) | 3.10 (2.4–3.80) | 5.57 (4.97–6.16) | 7.43 (6.50–8.36) |
| $b_3 * 10^{-4}$ | 0.93 (0.03–1.83) | 0.66 (0.02–1.30) | 0.39 (0.01–0.77) | 0.49 (0.01–0.97) |
| $\mu_{c_1}$ | 0.11 (0.06–0.16) | 0.13 (0.06–0.20) | 0.05 (0.03–0.07) | 0.30 (0.19–0.42) |
| $\mu_{c_2}$ | 0.044 (0.003–0.085) | 0.13 (0.03–0.22) | 0.22 (0.16–0.27) | 0.01 (.0004–0.02) |
| $\mu_{c_3}$ | 0.042 (0.002–0.083) | 0.29 (0.18–0.40) | 0.33 (0.25–0.42) | 0.32 (0.23–0.42) |
| $r$ | 0.06 (0.006–0.12) | 1.18 (0.83–1.54) | 1.01 (0.83–1.18) | 1.47 (1.10–1.83) |
| $\mu_{y_0} * 10^3$ | 0.681 (0.677–0.685) | 0.71 (0.35–1.07) | 0.68 (0.36–1.01) | 0.82 (0.46–1.18) |
| $x^*$ | 0.05 | 1.89 | 1.93 | 2.75 |
| $\sigma_{x_1}$ | 0.45 | 0.38 | 0.53 | 0.70 |
| $\sigma_{x_2}$ | 0.22 | 0.40 | 0.33 | 0.22 |
| $\sigma_{x_3}$ | 0.08 | 0.10 | 0.15 | 0.15 |
| $\sigma_y$ | 1.00 | 6.02 | 4.50 | 4.01 |

**Fig 5.**
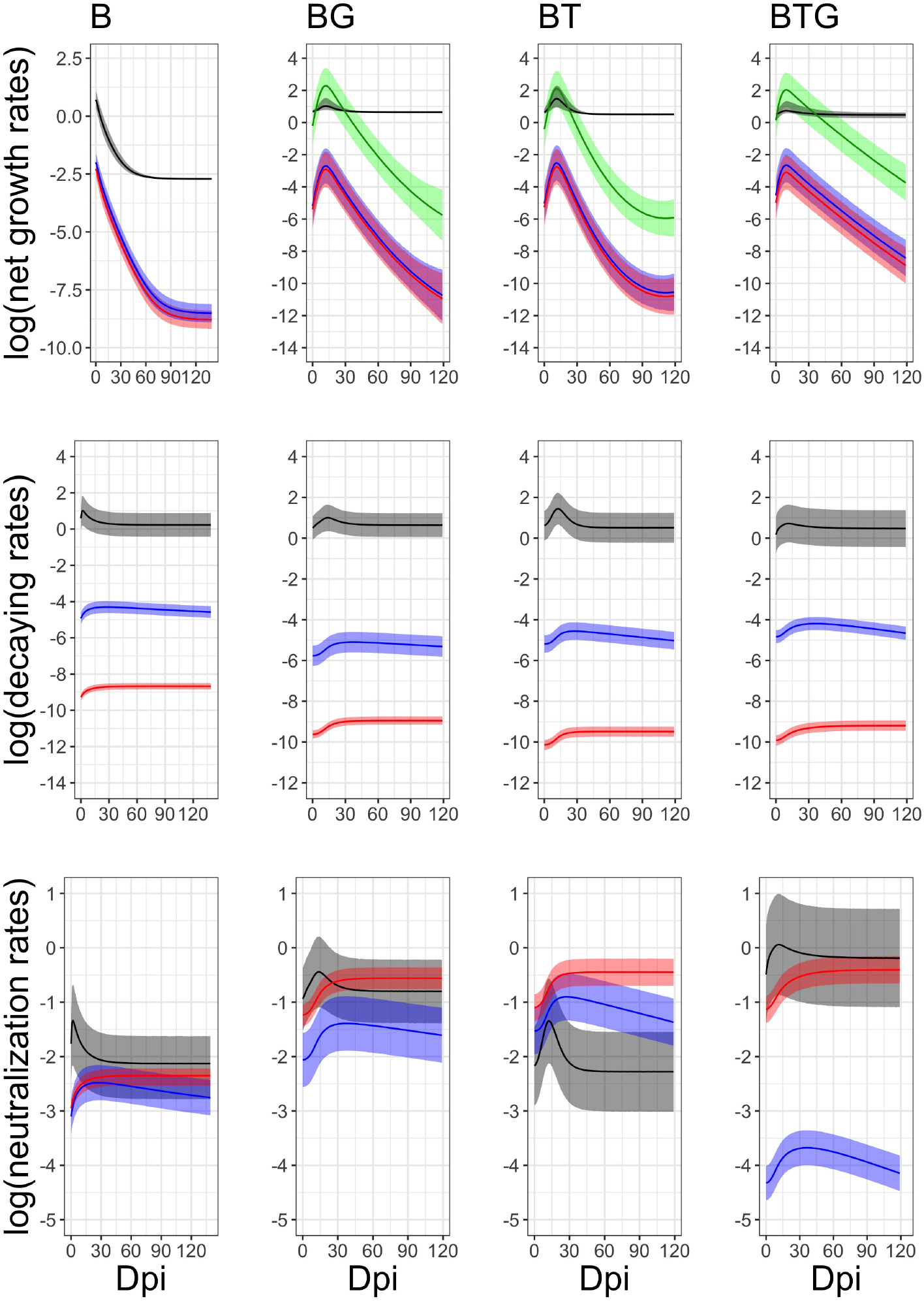
Simulated dynamics of the immune response in the blood and *B. bronchiseptica* infection in the whole respiratory tract by time, for B, BG, BT and BTG groups. Means (lines) and 95% CIs (shaded areas) are provided for neutrophils (black), IgA (blue), IgG (red) and *B. bronchiseptica* infection (green). Full details on the estimated rates, along with their credible intervals, are reported in Table 2.

Simulations highlighted clear differences between the four types of infection and the relative contribution of the three immune variables to *B. bronchiseptica* neutralization, *c*_1_, *c*_2_ and *c*_3_. Estimated parameters indicate that neutrophils primarily control *B. bronchiseptica* in the single infection, whereas a combination of neutrophils and specific IgG explains better the control of bacteria in the BTG rabbits (Table 2, Fig 5). IgG, followed by IgA, appears to play a more influential role in the BG and BT groups. It is important to note that the timing of these interactions is also critical. Given the fast reaction of neutrophils, they contributed to bacterial control early on in the infection, whereas specific IgG and, secondly, specific IgA were more important at a later time, as they decreased more slowly, like IgA, or remained consistently high over the course of the infection, as was the case for IgG (Fig 5).

### Modeling results: *B. bronchiseptica* infection explains variation in the dynamics of shedding between groups

For each individual rabbit, we examined the dynamics of shedding by linking the empirical shedding data to the estimated intensity of infection (see previous section for this estimation) through the zero inflated log-normal function and, from this, generated the simulated shedding data for each rabbit (details in Material and Methods). As previously noted, model fitting was performed on the whole dataset for the *B. bronchiseptica* only rabbits. Simulations showed that the rapid growth of *B. bronchiseptica* in the respiratory tract led to the rapid shedding of bacteria by host (Fig 6). The estimated time to reach the peak of shedding was between 10 and 12 days post infection, specifically: 9 days for BTG, 11 days for BT and 12 days for BG. For *B. bronchiseptica* only rabbits, the peak was right at the start of the trial, although this should be taken with caution, since data for the first 10 days of sampling were not available. Simulations also confirmed the consistently higher intensity of shedding in rabbits co-infected with helminths, the peak (log(CFU/day+1)) and 95% CIs were: 9.56 (2.49–14.63) for BTG, 8.97 (3.42–14.61) for BT, 9.02 (1.93–16.13) for BG and 6.52 (5.13–7.91) for *B. bronchiseptica* only rabbits, again, taking this latter result with caution. Among the three co-infections, simulations were able to capture the different shedding trends, albeit not statistically significant, as already noted for the experimental data. Rabbits from the BTG group exhibited the more rapid growth, the higher intensity of shedding and the slower decline post-peak, suggesting a synergistic effect of the two helminths. The BT group exhibited the most rapid decline, while rabbits from the BG group showed a trend intermediate between the BTG and BT groups (Fig 6).

**Fig 6.**
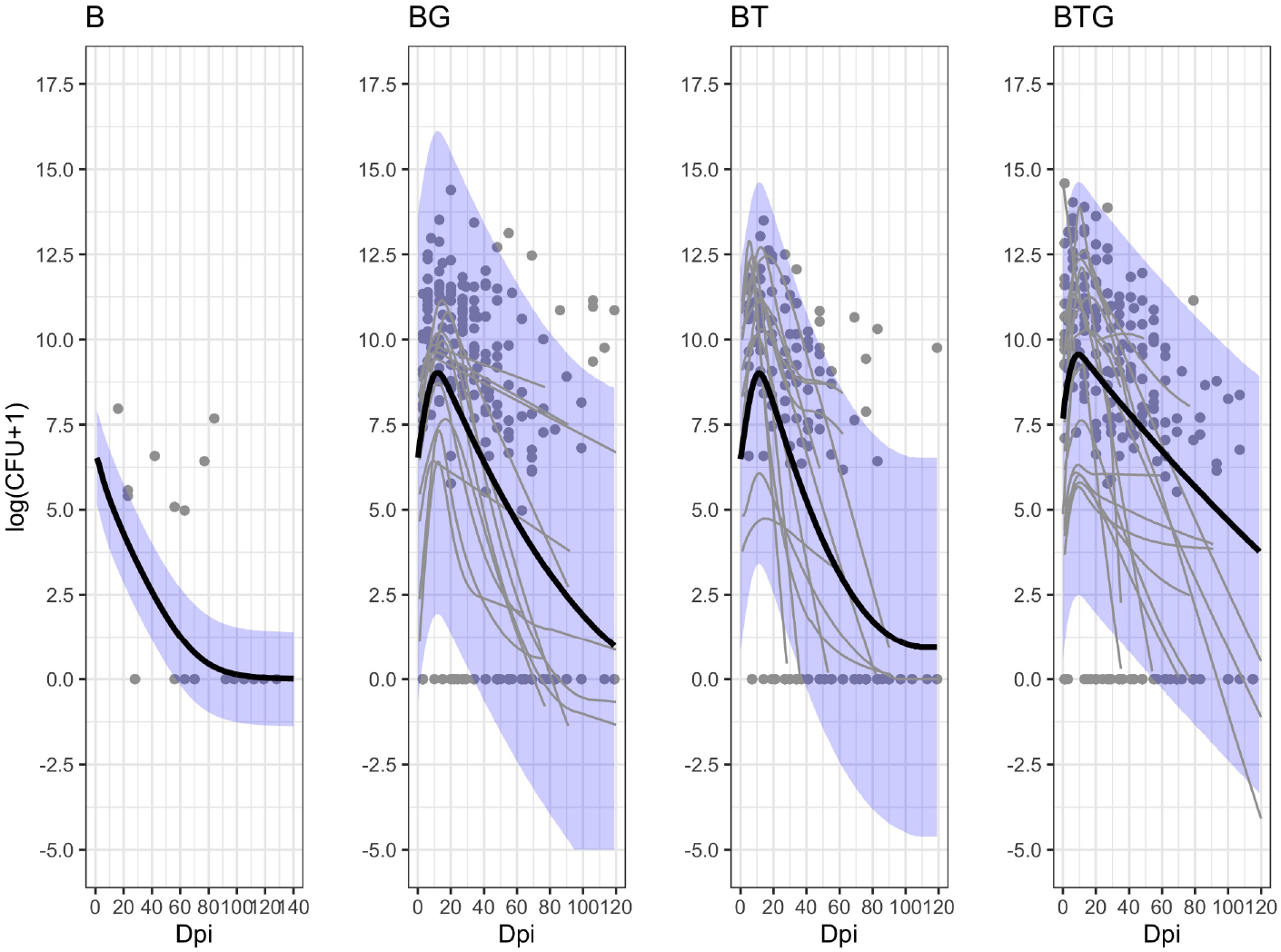
Estimated dynamics of *B. bronchiseptica* shedding for rabbits from the B, BG, BT and BTG groups. The empirical daily shedding events (gray points), the estimated individual trajectories (smoothed thin gray lines) and the estimated median group trends (smoothed thick black lines), with the related 95%CIs (blue areas), are presented for each group. Intensity of shedding is reported as total daily event.

At the host level, the rapid changes in the intensity of shedding between consecutive samplings, where bursts of bacteria alternate to low or no shedding in a matter of a few days, was weakly captured by model simulations. These rapid dynamics could not be explained by the intensity of infection and related immune responses we recorded. Instead, it is likely that alterations of the local conditions, such as variation in the severity of tissue inflammation and bacterial control, or in the amount of mucus formation and expulsion, had a stronger effect on these rapid changes. We also noted that co-infected rabbits sniffled more frequently than *B. bronchiseptica* only animals (Pathak’s pers. obs.), and, probably, further contributed to alter the frequency and intensity of shedding both between and within hosts.

### Sensitivity analysis

To explore in more detail how variation in shedding between types of infection was related to changes in key immune parameters, a sensitivity analysis was undertaken on model performance. Findings showed that an increase in the rate of bacterial neutralization *µ*_*c*_, and, secondly, in the immune growth rate *µ*_*a*_, led to a proportional decrease of the peak of shedding and, to a lesser extent, the time to reach this peak (Fig 7 and Fig 8). Among the three immune variables, the strongest negative impact on shedding was caused by changes in neutrophil (*c*_1_ or *a*_1_) and IgG (*c*_3_ or *a*_3_) rates. As expected, an increase in bacterial growth rate, *r*, was associated with a higher peak of shedding, and consequently a shorter time to reach this peak (Fig 7 and Fig 8). For a negative bacterial growth, namely, a decline in the intensity of infection, hosts carried on shedding if the decline was not too extreme to clear the infection. During strong negative growths (i.e. 75% or 50%), the peak of shedding and the zero time to reach this peak were represented by the initial inoculum.

**Fig 7.**
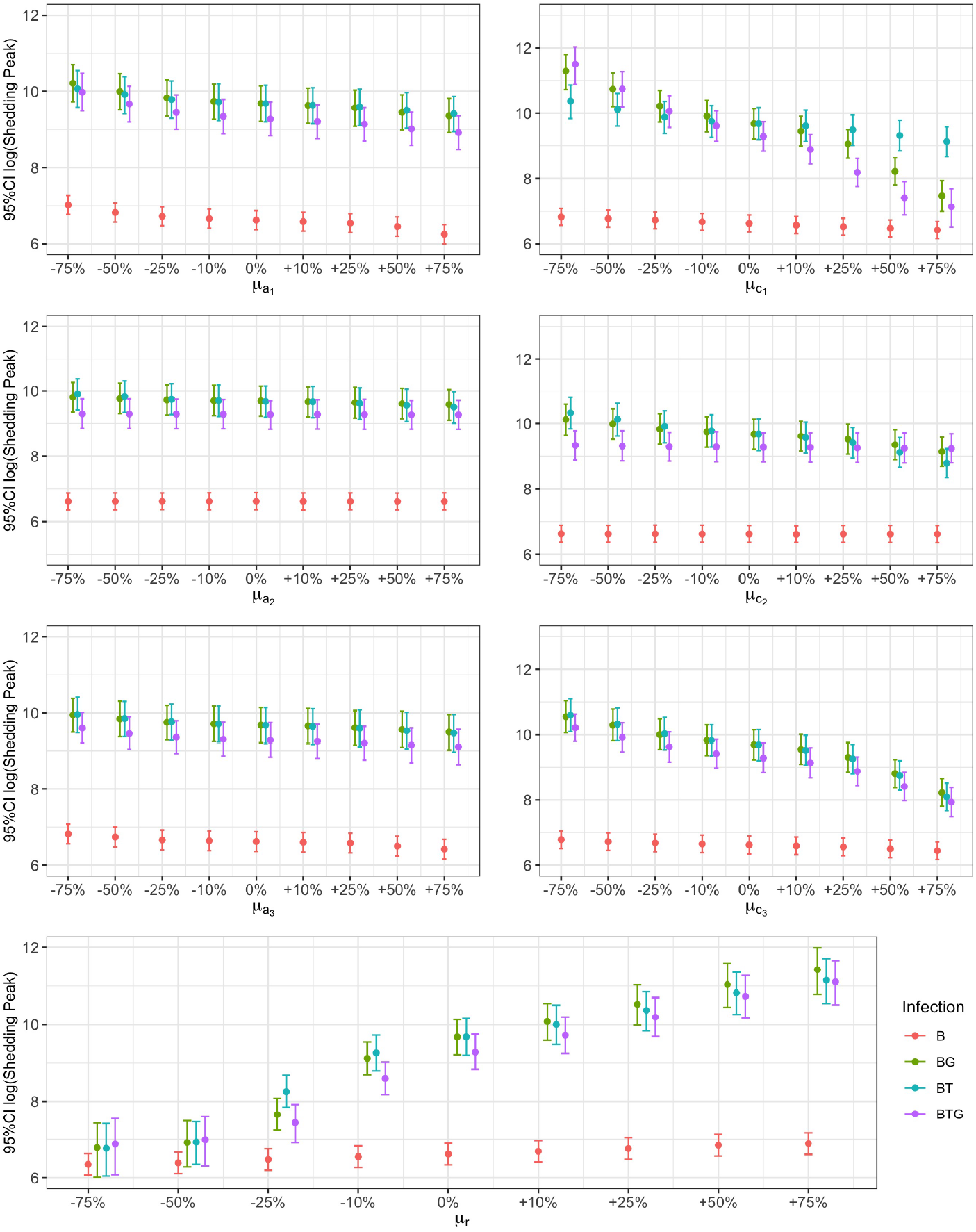
Relationship between *B. bronchiseptica* peak of shedding and percentile changes in neutrophils, IgA, and IgG. The growth *a*, and neutralizing *c*, rates of neutrophils (subscript 1), IgA (subscript 2) and IgG (subscript 3), including bacterial growth rate *r*, are reported by group. Mean estimates with 95% CIs are presented.

**Fig 8.**
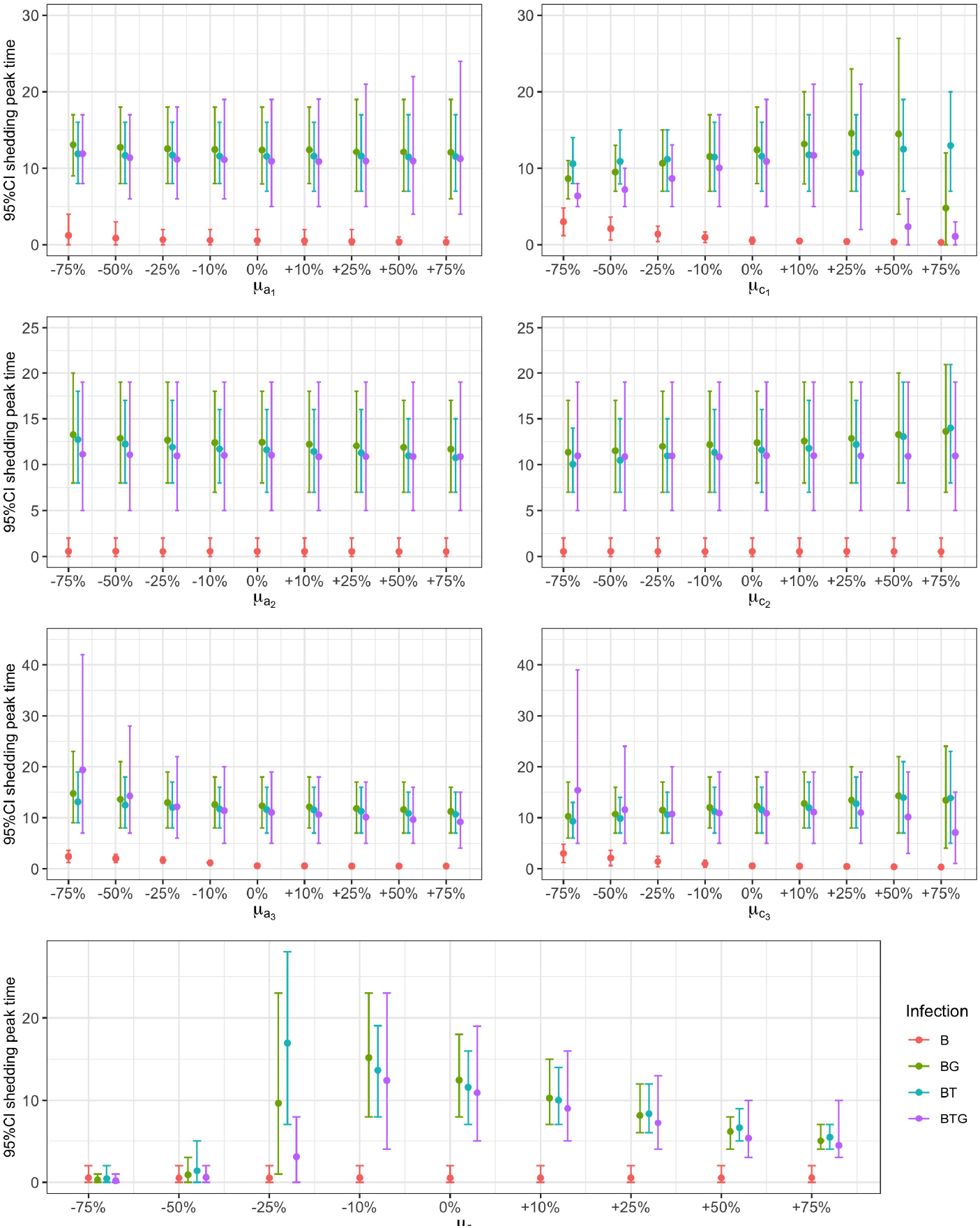
Relationship between *B. bronchiseptica* time to peak of shedding and percentile changes in neutrophils, IgA, and IgG. The growth *a*, and neutralizing *c*, rates of neutrophils (subscript 1), IgA (subscript 2) and IgG (subscript 3), including bacterial growth rate *r*, are reported by group. Mean estimates with 95% CIs are presented.

Overall, the three co-infections exhibited comparable trends and were more sensitive to immune changes than *B. bronchiseptica* only infected rabbits. The weak response of the latter group was caused by the very low bacterial growth rate (*r* =0.06) and thus shedding, indicative of a stronger immune control. These trends further confirm the evidence that helminths increase variation in the pattern of shedding both between and within groups. A strong deviation from the co-infected groups was observed for the BTG rabbits under an extremely high neutrophil control.

## Discussion

There is increasing evidence that gastrointestinal helminths can impact the severity and time course of respiratory bacterial infections [30, 35–38, 50], however, how they affect the dynamics of shedding remains to be determined. This is particularly important in regions where chronic helminthiases co-circulate with respiratory pathogens that are endemic or cause seasonal outbreaks. This is also relevant in areas where antimicrobial resistance is emerging as a threat, such that focusing on the treatment of parasitic helminths could be a rapid and effective way to reduce onward bacterial transmission. We investigated how two gastrointestinal-restricted helminth species, with contrasting infection dynamics, altered the shedding of the highly infectious *B. bronchiseptica* through the modulation of the immune response. Empirical findings showed that the two helminths, taken as a single species or as a pair, significantly increased the intensity, variation and duration of shedding as well as enhanced the frequency of supershedding events. Model simulations showed that these changes were related to the significant growth of bacteria in the respiratory tract. The prompted development of an immune response controlled the infection and led to the waning but not the complete cessation of shedding by the end of the experiments. Simulations revealed that changes in the relative contribution of neutrophils, IgA and IgG to bacterial neutralization explained differences in the pattern of shedding between types of infection. However, the rapid temporal changes observed at the individual level could not be explained by the infection–immunity relationship at the scale of analysis selected for this study.

We modeled the dynamics of shedding as representative of the infection in the whole respiratory tract, yet, simulations were able to capture the contrasting trends in the lungs and nasal cavity. The rapid decline of shedding in the first 90 days post infection can be explined by the immune-mediated clearance of *B. bronchiseptica* from the lungs, previously reported using data from these experiments [16, 20, 31], while the little but ongoing shedding later in the experiments can be the consequence of the weak control in the nasal cavity. Simulations showed that neutrophils quickly developed and contributed to bacteria neutralization early on, these were then followed by antibodies that developed more slowly and with an initial delay in the co-infected hosts. The two helminth species did not change the general trend of these immune variables but the timing and intensity of their relative contribution and, with this, the temporal changes in the intensity of shedding. We showed previously that both helminths elicit a strong type 2 immune response [34, 39, 40], however, while *T. retortaeformis* is cleared, or significantly reduced, from the small intestine, *G. strigosum* persists with high intensities in the stomach [33, 34, 39–41]. The faster decline of shedding in rabbits from the BT but not the BG group was caused by the stronger control of *B. bronchiseptica*, probably facilitated by the rapid clearance of *T. retortaeformis* [16]. As a consequence of this, rabbits from the BT group shed overall less and were characterized by a lower supershedding threshold (either at 99^*th*^ or 95^*th*^ percentile), compared to hosts from the BG group. The persistence of *G. strigosum* did not prevent bacteria clearance from the lower respiratory tract but reduced the decline of shedding and contributed to the higher number of supershedding events.

Previous studies have shown that serum antibodies (which include IgG isotypes and IgA) completely clear *B. bronchiseptica* from lungs and trachea of wild-type and B-cell-deficient mice within 3 days post inoculation [17]. Similarly, IgA was found to be necessary for the reduction of bacterial numbers in the nasal cavity of mice [18]. Our simulations showed that in the BT group, the specific IgG and IgA responses to bacterial neutralization developed more slowly but their impact was stronger than the neutrophil response, later in the trial. This neutralizing antibody response was also stronger in rabbits from the BT as opposed to the BG group. In this latter group, the low IgA reaction and the slow development of IgG, albeit with strong neutralizing properties from day 30 post infection, contributed to the decline of shedding but not as fast as rabbits from the BT group. Taken together, these findings suggest that antibodies were critical for the rapid decline of *B. bronchiseptica* shedding in the BT rabbits and the slower reduction in the BG group.

Our model assumes that neutrophils are stimulated by, and directed against, *B. bronchiseptica* replication in the respiratory tract. However, neutrophils are also produced during helminth establishment, probably as a consequence of gut bacteria infiltration in the helminth-damaged mucosa [16, 39, 40]. By driving the additional recruitment of neutrophils, *T. retortaeformis* could have facilitated *B. bronchiseptica* clearance from the lungs and the rapid reduction of shedding in rabbits from the BT group. While neutrophils are also produced during *G. strigosum* infection [39], the lack of parasite clearance and the constant stimulation of a type 2 response against this helminth, could explain the extended shedding of rabbits from the BG group. The evidence that gut-restricted helminths enhance the neutrophil response to respiratory bacteria was further confirmed in mice co-infected with *Pseudomonas aeruginosa* and *Heligmosomoides polygyrus* [38]. This study also reported an increase of CD4+ T cells and Th2 cytokine expression in the lungs; a mechanism that, perhaps, could also contribute to explain the higher shedding reported for the group of BG rabbits. These findings contrast to some extent with the work by Rolin et al [19] using TLR4-mutant mice infected with *B. bronchiseptica* alone. These mice showed a poor control of bacterial growth and shedding, and were characterized by a heavy neutrophil infiltration in the lungs. To note is that the depletion of neutrophils did not affect the level of infection but decreased individual shedding.

Interestingly, in the BTG triple infection, the two helminths appeared to have a synergistic effect, in that they increased the level of shedding, reduced its rate of decline and increased the probability of supershedding events. Simulations suggest that, compared to dual infections, these trends were associated with the substantially reduced neutralizing properties of IgA, and less so of IgG. As previous commented for dual infections, we suggest that a robust type 2 bias environment was probably present in the respiratory tract of these rabbits and further contributed to enhance the intensity and duration of shedding when both helminths were present. Our previous studies of a natural population of rabbits showed that 65% of the hosts were co-infected with *B. bronchiseptica* and *G. strigosum*, and both bacterial prevalence and helminth intensity increased with rabbit age [20]. Although we did not examine these patterns in dual or triple infections with *T. retortaeformis*, we found that about 60% of adult rabbits were BTG triple infected (unpubl. data). Given that *T. retortaeformis* and *G. strigosum* are commonly found in natural populations of rabbits, and considering that helminth-infected hosts represent a high percentage in these natural populations, it is not surprising if these helminths play an important role to the rapid transmission of *B. bronchiseptica* and persistence in natural rabbit systems. We can also speculate that a similar pattern, namely, increased bacterial shedding and enhanced transmission, should be expected in humans co-infected with gastrointestinal helminths and *B. pertussis* or *B. parapertussis*.

The *B. bronchiseptica* group was characterized by a low level of shedding, a high frequency of null events and few supershedding. Simulations indicated that these trends were associated with a prompt and relatively strong response of neutrophils followed by a much lower antibody reaction. Previous Boolean modeling of the whole immune network to *B. bronchiseptica* infection in the lungs, indicated that the antigen-antibody complexes, at which IgA and IgG contribute, and phagocytes/ phagocytosis, activated by neutrophils among others, were important for bacterial removal [16]. Our simulations, using a different modeling approach, add to this by suggesting that the early and fast bacterial neutralization by neutrophils, and secondly antibodies, could have set host immunity to better control bacterial growth and, with this, shedding. As confirmed by the sensitivity analysis, percentile changes of the already low growth rate had negligible effects on the peak of shedding or the time to reach the peak.

A common feature among rabbits, irrespective of the infection group they belong to, was the rapid fluctuation in the level of bacteria shed over time, including the unpredicted supershedding events. We could not explain these rapid changes based on our scale of analysis, relaying on measurements of the immune response from blood and intensity of infection from the whole respiratory tract. Most likely, other processes generated the bursts and timing of (super)shedding observed. These could include, but are not limited to, immune-mediated control of bacteria turnover at the epithelium/ mucosa level or stochastic events during bacteria release, such as random mucus discharge, and also variation in host behavior. Overall, our findings are consistent with other systems where fast and intermittent shedding has been reported both from single [6, 19, 42–44] and co-infected hosts [12, 45]. Our model framework and variables selected, prevented us from capturing the intrinsic and local mechanism responsible for these rapid changes, including the processes generating supershedding events, and more work at the epithelial level is need to explain these patterns and the relative contribution of the lower and upper respiratory tract.

As illustrated by our findings, the definition of supershedder can become problematic when hosts are infected with more than one parasite/pathogen. Our analysis showed that a cut-off value based on single infection can lead to the overestimation of supershedding in co-infected hosts. By using a percentile threshold for each infection, we can provide a more accurate estimation but this implies different cut-offs, which could be difficult to compare if species are phylogenetically different. We used two closely related helminth species and if we consider the 95^*th*^ percentile thershold for each type of infection we found that between 20% and 40% of the co-infected rabbits generated at least one supershedding event by contact. This is a significant percentage in our infected groups, especially if about 65% of shedding led to null events. It is also important to note that the intensity of these events is significantly higher compared to rabbits with bacteria only, and thus the risk of onwards transmission, given a contact, exponentially increases for co-infected hosts.

This study adds novel insights into the role of gastrointestinal helminths to the dynamics of *B. bronchiseptica* shedding. We used a parsimonious mechanism of immune regulation based on previous work by others and ourselves. While we focused on antibodies and neutrophils our model framework can be adapted to include the impact of other immune variables, such as components of the innate immune response or relationships between immune variables. To reduce model complexity, we did not explicitly quantify the dynamics of infection of the two helminths although this can be explored in future work, including how bacteria affect the dynamics and shedding of helminths.

Given that one quarter of the global human population is infected with helminths and considering that respiratory infections are among the top ten causes of death by infectious diseases worldwide, understanding the modulatory role of helminth species to respiratory infections is important for developing treatments targeted to specific co-infection settings. The ability to detect and, ideally, control the high shedders and/or supershedders is also critical for reducing the risk of disease outbreak and spread, and should not be overlooked any longer.

## Materials and Methods

### Ethic Statement

Animals were housed in individual cages with food and water *ad libitum* and a 12h day/night cycle, in compliance with Animal Welfare Act regulations as well as the Guide for the Care and Use of Laboratory Animals. All animal procedures, including infections with the bacterium and the two helminth species, weekly blood collection and pathogen/parasite sampling at fixed time, were approved by the Institutional Animal Care and Use Committee of The Pennsylvania State University (IACUC 26082). All animal work complied with guidelines as reported in the Guide for the Care and Use of Laboratory Animals. 8th ed. National Research Council of the National Academies, National Academies Press Washington DC.

### Bacteria strain and culture

We used B. bronchiseptica strain RB50 for all experiments. Bacteria were grown on Bordet-Gengou (BG) agar supplemented with 10% defibrinated sheep blood and streptomycin (20 µg/ml). The inoculum was prepared by growing the bacteria in Stainer-Scholte (SS) liquid culture medium at 37°C overnight. For the infection, bacteria were re-suspended in sterile phosphate-buffered saline (PBS) at a density of 5 *∗* 10^4^ CFUs/ml, which was confirmed by plating serial dilutions of the inoculum on BG blood agar plates in triplicate [20].

### Laboratory Infections

*B. bronchiseptica* single infection (B) and dual infections with either *T. retortaeformis* (BT) or *G. strigosum* (BG) are described in detail in Pathak et al. [20, 31] and Thakar et al. [16]. The triple infection (BTG) followed the same experimental design and laboratory procedures of the dual infections. A general description of the components relevant to the current study is reported here. New Zealand White, two months old, male rabbits (Harlan, USA), were simultaneouly challenged with *B. bronchiseptica* and helminths as follow: i-*B. bronchiseptica* (B) single infection: 32 infected and 16 controls, ii-*B. bronchiseptica*-*Graphidium strigosum* (BG): 31 infected and 16 controls, iii-*B. bronchiseptica*-*T. retortaeformis* (BT): 32 infected and 16 controls, and iv-the three agents together (BTG): 32 infected and 16 controls. Infection was performed by pipetting in each nare 2.5 * 10^4^CFU/ml of bacteria diluted in 0.5 ml of PBS. For the co-infections, animals also received, simultaneously by gavage, a single inoculum of water (5 ml) with either 5500 *T. retortaeformis* or 650 *G. strigosum* third stage infective larvae, or both. Helminth doses followed natural infections [32, 33]. Control animals were scam inoculated with 1 ml of sterile PBS in the nares and gavaged with 5ml of water. The dynamics of infection, shedding and related immune response were then followed for 120 days (150 days for B) by sacrificing four infected and two control animals at fixed days post infection, as follow, B: 3, 7, 15, 30, 59, 90, 120, 150 days; BG: 7, 14, 30, 44, 62, 76, 90, 120 days; BT: 5, 8, 15, 31, 47, 61, 91, 120 days; BTG 7, 14, 30, 45, 60, 75, 90, 120 days. Data were collected both for *B. bronchiseptica* and the helminths, including the host immune response.

### Bacteria shed enumeration

At the start of each infection, a subset of infected rabbits was randomly selected (i.e. B=16, BG=23, BT=20 and BTG=24) to quantify the intensity of bacteria shed over time and by contact with a BG-blood agar petri dish [20]. Shedding by contact with a surface exemplifies the natural transmission of a respiratory pathogen without disruption of the bacteria population through swabbing. Shedding was assessed once a week for B single infection and every 2 (BT) to 3 (BG and BTG) times a week for the co-infected rabbits. The choice to use a different number of hosts and a different frequency of sampling was determined by logistical reasons. Technical difficulties prevented the collection of bacteria in the first 10 days post-infection for *B. bronchiseptica* only rabbits. At every event, rabbits were allowed to play with the petri dish by direct oral-nasal contact and for a maximum of 10 minutes. Plates were removed earlier if animals chewed the plastic or the agar, and the duration of each interaction was recorded. Plates were then incubated at 37°C for 48 hours and colonies counted and scaled to the interaction time (CFUs/s). If there was an interaction but plates resulted negative, shedding was considered to be zero; the lack of interaction (animals not interested in the plate) was recorded as a missing point.

### Systemic immune response

As representative of the immune response to *B. bronchiseptica*, we selected neutrophils and species-specific antibodies IgA and IgG. Previous studies showed that these three variables are required to clear *B. bronchiseptica* from the lungs and to reduce the colonization in the nasal cavity [14, 16–18, 46]. For example, the modeling of the immune network in the lungs of mice and rabbits showed that the lack of antibody production, by B cells deletion, prevented bacterial clearance [14, 16]. Consistent with these findings, experimental studies showed that adoptive transfer of serum antibodies in mice led to the removal of *B. bronchiseptica* by day 3 post inoculation [17]. Similarly, simulations showed that peripheral neutrophils, recruited via pro-inflammatory cytokines, contributed to the activation of phagocytic cells, which are fundamental for bacterial neutralization [14, 16].

Methodologies applied to quantify the three immune variables are described Pathak et al. [20, 31] and Thakar et al. [16]. Briefly, for every rabbit blood was collected once a week for neutrophils or twice a week for antibodies. Neutrophil concentration was measured using whole blood (0.2 ml) stored in EDTA (Sartorius, Germany) and analyzed using Hemavet-3 hematology system (Drew Scientific, USA). Species-specific IgA and IgG were estimated from blood serum samples using *B. bronchiseptica* as a source of antigen and ELISA [20]. Measurements were performed in duplicates with all plates having high, low and background controls. Values were expressed as immunosorbent Optical Densities (OD) and then standardized into Optical Density index [39]. Plate preparation and dilutions, including the preparation of high (strongly reacting animals) and low (non-reacting animals from prior to the infection) pools and checkerboard titrations, are detailed elsewhere [16, 20, 31]. For the triple infection, antibody quantification and dilutions followed Pathak et al. [20]. The initial baseline immune conditions of every rabbit were available from data collected the week before the infection, if data were missing we used the average value from control rabbits. The neutrophil equilibrium level, was available as the average value from control rabbits sampled throughout the experiments, specifically cells/ml B= 1.323, BG= 2.730, BT= 1.819 and BTG= 2.289.

### Statistical analysis

To investigate the degree of variation in the intensity of shedding within each group, and identify the supershedding events associated with this variation, the frequency of shedding events was examined. Shedding events were grouped into discrete intervals based on their intensity and the frequency calculated for each interval. Supershedding events were identified as the cases above the 99^*th*^ (or 95^*th*^) percentile estimated for each infection. As a comparison, we also considered the 99^*th*^ percentile of *B. bronchiseptica* single infection as a reference threshold for all the infections, and estimated the number of supershedding cases above this common cut-off value.

### Model framework

The within-host mechanisms that affect the dynamics of *B. bronchiseptica* shedding were examined for each individual rabbit in two steps. First, we developed a dynamical model that describes the dynamics of neutrophils, specific IgA and IgG, and their interaction with the bacterial infection in the whole respiratory tract. Second, a Bayesian approach was then used to link this dynamical model to the empirical longitudinal data by: i-fitting the model to experimental time series of neutrophils, IgA, and IgG and ii-fitting the intensity of *B. bronchiseptica* infection, estimated from the dynamical model, to the experimental shedding data. Below we describe the dynamical framework and in the next section we report on how this model was fitted to the empirical data, including how parameter calibration was performed.

### Dynamical model

The time-dependent interactions between immune variables and *B. bronchiseptica* infection is described by the following system of ordinary differential equations:

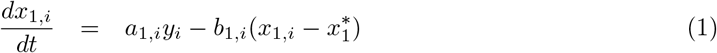

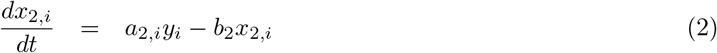

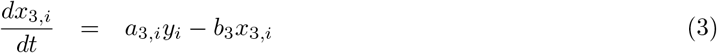

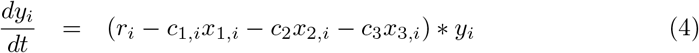

where *x*_1,*i*_(*t*), *x*_2,*i*_(*t*) and *x*_3,*i*_(*t*) are neutrophils, IgA, and IgG, respectively, of rabbit *i*^*th*^ at time *t. y*_*i*_(*t*) is the intensity of bacteria in the respiratory tract of this same rabbit at time *t*. The parameters *a*_1,*i*_, *b*_1,*i*_ and *c*_1,*i*_ describe the rate of growth, decay and bacteria clearance, respectively, for neutrophils in rabbit *i*^*th*^. Similarly, *a*_2,*i*_, *b*_2_ and *c*_2,*i*_ describe the per capita rates of growth, decay and baterial clearance for IgA, while *a*_3,*i*_, *b*_3_ and *c*_3,*i*_ are the rates representing IgG (Table 1). The baseline immune conditions of the host before the infection were represented by *x*_1,*i*_(0), *x*_2,*i*_(0) and *x*_3,*i*_(0), while 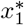 describes the equilibrium level of neutrophils post infection; this was not included for IgA or IgG since the time to reach equilibrium extended beyond the course of the experiments. *r*_*i*_ represents the per capita bacterial replication rate. The full description of model parameters and variables is reported in (Table 1).

We assumed that the three immune variables proportionately increase with the intensity of infection, *y*_*i*_(*t*), and neutralize *B. bronchiseptica* following a Lotka-Volterra type of relationship [47–49]. We also assumed that helminths affect *B. bronchiseptica* infection, and thus shedding, by altering the magnitude and time course of the neutrophil, IgA and IgG response.

### Observation Model

We applied the dynamical model described above to every rabbit of the four infections, independently. Individual parameters could vary among hosts as independent samples from a joint log-normal distribution, while parameters that were shared among rabbits within the same group, i.e. *b*_2_ and *b*_3_, were kept constant. This hierarchical set-up allowed us to have individual responses that varied in term of amplitude, time to peak and decay rate, while keeping these curves from deviating too much from each other. For a given rabbit *i*^*th*^ at time *t*, every empirical immune variable, log-transformed, was assumed to follow a normal distribution with mean, *µ*, and variance, *σ*, as follow:

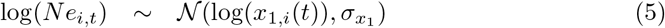

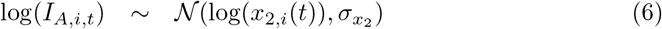

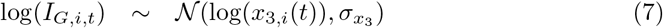

The empirical amount of *B. bronchiseptica* shed by rabbit *i*^*th*^ at time *t*, is directly proportional to the level of bacterial infection *y*_*i*_(*t*). This formulation makes the assumption that shedding is representative of the intensity of infection in the whole respiratory tract. Also, the probability of having a shedding event is independent both of time since innoculation, i.e. a shedding can occur anytime during the experiment, and the time of interaction with the petri dish, i.e. shedding can occur any time during this interaction. Shedding was then related to the dynamics of infection via a zero-inflated log-normal relationship, to account for the high fraction of zero-shedding events (Fig 3), as follow:

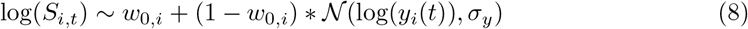

where *w*_0,*i*_ is the fraction of zero shedding events recorded for a given host. The estimated shedding was scaled up and quantified as total amount of bacteria shed by a rabbit in a day.

### Model selection and parameter calibration

The dynamical model described in equations (1)-(4) included parameters that were shared among rabbits within the same group, specifically:

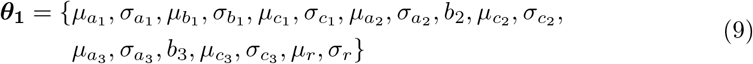

where *µ* and *σ* are the means and standard deviations, respectively, of the corresponding normal distributions of the parameters *a, b, c* and *r* for each group. Likewise, the set of parameters ***θ***_**2**,***i***_ represents the estimates of every individual rabbit within a group, as:

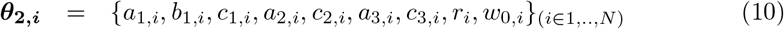

where N is the number of infected rabbits. The resulting hyper prior distribution for rabbit *i*^*th*^ is:

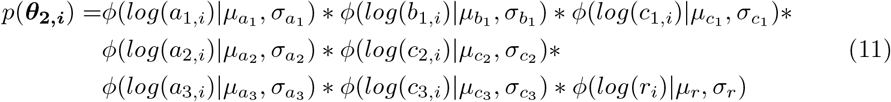

The resulting joint-likelihood function for the observed longitudinal time-series of all infected rabbits is:

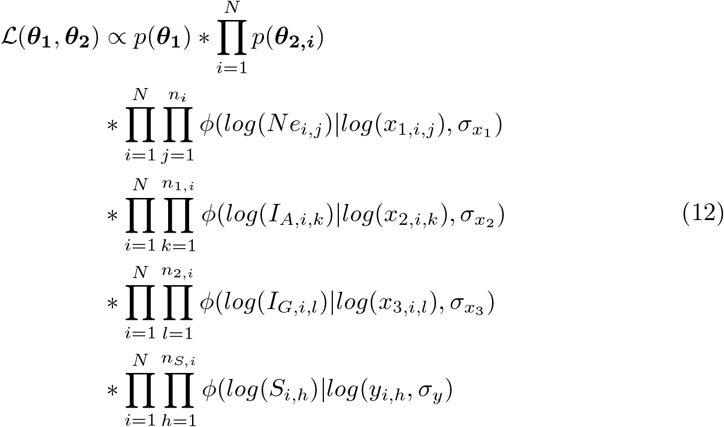

where *n*_*i*_, *n*_1,*i*_, *n*_2,*i*_, and *n*_*S,i*_ are the number of measurements for neutrophil, IgA, IgG and bacteria shed, respectively, for rabbit *i*^*th*^. The four subscripts *j, k, l, h* are the time index for neutrophil, IgA, IgG, and bacteria shed, respectively. *Ne*_*i,j*_, *I*_*A,i,k*_, *I*_*G,i,l*_, and *S*_*i,h*_ are the neutrophils, IgA, IgG and shedding measurements at the respective time points for rabbit *i*^*th*^. The notation *φ*(*x*|*µ, σ*^2^) represents the normal probability density of *x* with mean *µ* and variance *σ*^2^. Since there is no knowledge on the values of ***θ***_**1**_ we used a weakly normal prior for *p*(***θ***_**1**_).

Bayesian model simulations were performed using Halmiltonian Monte Carlo (HMC) algorithm in Stan package (version 2.18.0). Four chains were run, each with 100,000 iterations with 40,000 burn-in iterations. Parameter estimates were diagnosed to make sure that the chains were well-mixed with no divergent transitions post-burning.

Simulations were one-day time step increment and parameters were scaled to account for differences in their relative magnitude. Rabbits that had less than four shedding data points were excluded from the modeling, specificaly we used: 17 BT, 22 BG and 23 BTG rabbits. For the *B. bronchiseptica* group, out of 16 rabbits sampled only 7 could be used for the modeling, however, given this low number of hosts and some difficulties with model convergence, the modeling was carried out using the whole group data and results were presented for the entire group.

Finally, to improve model performance, two different assumptions were examined for the quantification of IgA and IgG decay rates: i-the rates are assumed to vary between individuals within the same infection group (full model), and ii-the rates are assumed to be the same for all rabbits within the same infection group (reduced model). This was motivated by the observation that IgA and IgG decay rates were relatively constant amongs rabbits within the same group. The reduced model was chosen as a balance between the number of fitted parameters (difference between full and reduced model in parameter number Δ: 47 for BG, 45 for BT and 49 for BTG) and model performance (Δ*BIC*: -80.5 for BG, -75.2 for BT and -90.5 for BTG).

### Sensitivity analysis

To examine how key immune parameters affected the magnitude and timing of the peak of shedding, a sensitivity analysis was performed. As representative of shedding we considered the peak of shedding and the time to reach the peak, including bacteria growth rate *r*, while for the three immune variables we selected the rate of bacterial neutralization *µ*_*c*_ and the growth rate *µ*_*a*_ of each population of cells or proteins. Analyses were carried out using the mean parameters of each type of infection as explained in Table 1. Specifically, for either *c* or *a* the estimated optimal value presented in Table 2 was changed by an incremental percentage, while keeping all the other parameters fixed, and the consequent changes in the amplitude and timing of the shedding peak quantified. The same exercise was repeated using incremental changes of *r*. For each infection group 1000 simulations were run for each of the seven parameters and we reported the mean values and the 95% confident interval from these 1000 simulations.

## Acknowledgments

The authors would like to thank Kathleen Creppage and Chad Pelensky for their research assistance during animal work and sample processing. This study, IMC and AKP were supported by Human Frontier Science Program (RGP0020/2007-C) and National Science Foundation-DEB (1145697). The funders had no role in study design, data collection and analysis, decision to publish or preparation of the manuscript.

## Supporting Information captions

**SI-1. Variation in neutrophil, IgA and IgG responses to *B. bronchiseptica* infection in B, BG, BT and BTG**.

Here, we report on experimental and simulated results for the three immune variables, presented as individual host data or group means.

**SI-2. Model parameter estimates from posterior distribution in BG, BT and BTG**.

The individual and group averages are reported for the immune parameters and bacterial infection.

## References

1. Keeling MJ, Woolhouse ME, Shaw DJ, Matthews L, Chase-Topping M, et al. (2014) Dynamics of the 2001 UK foot and mouth epidemic: stochastic dispersal in a heterogeneous landscape Science 294: 813–817.

2. Lloyd-Smith JO, Schreiber SJ, Kopp PE, Getz WM (2005) Superspreading and the effect of individual variation on disease emergence. Nature 438: 355–359.

3. Chase-Topping M, Gally D, Low C, Matthews L, Woolhouse M (2008) Super-shedding and the link between human infection and livestock carriage of Escherichia coli O157. Nature Rev Microb 6: 904.

4. Gopinath S, Hotson A, Johns J, Nolan G, Monack D (2013) The systemic immune state of super-shedder mice is characterized by a unique neutrophil-dependent blunting of TH1 responses. PLoS Pathog 9: pe1003408.

5. Chen WJ, Yang JY, Lin JH, Fann CS, Osyetrov V, et al. (2006) Nasopharyngeal shedding of Severe Acute Respiratory Syndrome—associated coronavirus is associated with genetic polymorphisms. Clin Inf Dis 42: 1561–1569.

6. Hadinoto V, Shapiro M, Sun CC, Thorley-Lawson DA (2009) The dynamics of EBV shedding implicate a central role for epithelial cells in amplifying viral output. PLoS Pathog 5: e1000496.

7. Leung NH, Xu C, Ip DK, Cowling BJ (2015) The fraction of influenza virus infections that are asymptomatic: a systematic review and meta-analysis. Epidemiology 26: 862.

8. Sheth PM, Danesh A, Sheung A, Rebbapragada A, Shahabi K, et al. (2006) Disproportionately high semen shedding of HIV is associated with compartmentalized cytomegalovirus reactivation. J Inf Dis 193: 45–48.

9. Graham AL, Cattadori IM, Lloyd-Smith JO, Ferrari MJ, Bjørnstad ON (2007) Transmission consequences of coinfection: cytokines writ large?. Trends Parasit 23: 284–291.

10. Richard AL, Siegel SJ, Erikson J, Weiser JN (2014) TLR2 signaling decreases transmission of Streptococcus pneumoniae by limiting bacterial shedding in an infant mouse Influenza A co-infection model. PLoS Pathog 10: p.e1004339.

11. Smith AM, Adler FR, Ribeiro RM, Gutenkunst RN, McAuley JL, et al. (2013) Kinetics of coinfection with influenza A virus and Streptococcus pneumoniae. PLoS Pathog 9: p.e1003238.

12. Byrne CM, Johnston C, Orem J, Okuku F, Huang ML, et al. (2019) Increased oral Epstein-Barr virus shedding with HIV-1 co-infection is due to a combination of B cell activation and impaired cellular immune control. bioRxiv 587063.

13. Goodnow RA (1980) Biology of Bordetella bronchiseptica. Microb Rev 44: 722–738.

14. Thakar J, Pilione M, Kirimanjeswara G, Harvill ET, Albert R (2007) Modeling systems-level regulation of host immune responses. PLoS Comput Biol 3: e109.

15. Thakar J, Saadatpour-Moghaddam A, Harvill ET, Albert R. (2009) Constraint-based network model of pathogen–immune system interactions. J R S Interface 6: 599–612.

16. Thakar J, Pathak AK, Murphy L, Albert R, Cattadori IM (2012) Network model of immune responses reveals key effectors to single and co-infection dynamics by a respiratory bacterium and a gastrointestinal helminth. PLoS Comput Biol 8: e1002345.

17. Kirimanjeswara GS, Mann PB, Harvill ET (2003) Role of antibodies in immunity to Bordetella infections. Infect Immun 71: 1719—1724.

18. Wolfe DN, Kirimanjeswara GS, Goebel EM, Harvill ET (2007) Comparative role of immunoglobulin A in protective immunity against the Bordetellae. Infect Immun 75: 4416–4422.

19. Rolin O, Smallridge W, Henry M, Goodfield L, Place D, Harvill ET (2014) Toll-like receptor 4 limits transmission of Bordetella bronchiseptica. PLoSOne 9: e85229

20. Pathak AK, Creppage KE, Werner JR, Cattadori IM (2010) Immune regulation of a chronic bacteria infection and consequences for pathogen transmission. BMC Microb 10: 1–9.

21. Long GH, Sinha D, Read AF, Pritt S, Kline B, et al. (2010) Identifying the age cohort responsible for transmission in a natural outbreak of Bordetella bronchiseptica. PLoS Pathog 12: p.e1001224.

22. Woolfrey BF, Moody JA (1991) Human infections associated with Bordetella bronchiseptica. Clin Microb Rev 4: 243–255.

23. de Cellés MD, Magpantay FM, King AA, Rohani P (2018) The impact of past vaccination coverage and immunity on pertussis resurgence. Science Transl Med 10(434).

24. Center for Disease Control and Prevention (2021) https://www.cdcgov/pertussis/countries/indexhtml. Opened: Jan, 6, 2012.

25. Brockmeier SL, Loving CL, Nicholson TL, Palmer MV (2008) Coinfection of pigs with porcine respiratory coronavirus and Bordetella bronchiseptica. Vet Microbiol 128: 36—47.

26. Loving CL, Brockmeier SL, Vincent AL, Palmer MV, Sacco RE, et al. (2010) Influenza virus coinfection with Bordetella bronchiseptica enhances bacterial colonization and host responses exacerbating pulmonary lesions. Microb Pathog 49: 237—245.

27. Schulz BS, Kurz S, Weber K, Balzer HJ, Hartmann K (2014) Detection of respiratory viruses and Bordetella bronchiseptica in dogs with acute respiratory tract infections. Vet J 201: 365–369.

28. Kureljušić B, Weissenbacher-Lang C, Nedorost N, Stixenberger D, Weissenböck H (2016) Association between Pneumocystis spp and co-infections with Bordetella bronchiseptica, Mycoplasma hyopneumoniae and Pasteurella multocida in Austrian pigs with pneumonia. Vet J 207: 177–179.

29. Hughes HR, Brockmeier SL, Loving CL (2018) Bordetella bronchiseptica colonization limits efficacy, but not immunogenicity, of live-attenuated influenza virus vaccine and enhances pathogenesis after influenza challenge. Front Imm 9: 2255.

30. Brady, O’Neill SM, Dalton JP, Mills KH (1999) Fasciola hepatica suppresses a protective Th1 response against Bordetella pertussis. Inf Immun 67: 5372–5378.

31. Pathak AK, Pelensky C, Boag B, Cattadori IM (2012) Immuno-epidemiology of chronic bacterial and helminth co-infections: observations from the field and evidence from the laboratory. Int J Parasit 42: 647–655.

32. Cattadori IM, Boag B, Bjørnstad ON, Cornell SJ, Hudson PJ (2005) Peak shift and epidemiology in a seasonal host–nematode system. Proc R Soc B 272: 1163–1169.

33. Cattadori, IM, Boag, B, Hudson PJ (2008) Parasite co-infection and interaction as drivers of host heterogeneity. Int J Paras 38: 371–380.

34. Cattadori IM, Pathak AK, Ferrari MJ (2019) External disturbances impact helminth–host interactions by affecting dynamics of infection, parasite traits, and host immune responses. Ecol Evol 9: 13495–13505.

35. Lass S, Hudson PJ, Thakar J, Saric J, Harvill E, et al. (2013) Generating super-shedders: co-infection increases bacterial load and egg production of a gastrointestinal helminth. J R S Interface 10: p20120588.

36. Ezenwa VO, Etienne RS, Luikart G, Beja-Pereira A, Jolles AE (2010) Hidden consequences of living in a wormy world: nematode-induced immune suppression facilitates tuberculosis invasion in African buffalo. Am Nat 176: 613–624.

37. Babu S, Nutman TB (2016) Helminth-tuberculosis co-infection: an immunologic perspective. Trends Immunol 37: 597–607.

38. Long SR, Lanter BB, Pazos MA, Mou H, Barrios J, et al. (2019) Intestinal helminth infection enhances bacteria-induced recruitment of neutrophils to the airspace. Scie Rep 9: 1–13.

39. Murphy L, Nalpas N, Stear M, Cattadori IM (20) Explaining patterns of infection in free-living populations using laboratory immune experiments. Parasit Immunol 33: 287–302.

40. Murphy L, Pathak AK, Cattadori IM (2013) A coinfection with two gastrointestinal nematodes alters host immune responses and only partially parasite dynamics. Parasit Immunol 35: 421–432.

41. Mignatti A, Boag B, Cattadori IM (2016) Host immunity shapes the impact of climate changes on the dynamics of parasite infections. PNAS 113: 2970–2975.

42. Schiffer JT, Abu-Raddad L, Mark KE, Zhu J, Selke S, et al. (2009) Frequent release of low amounts of herpes simplex virus from neurons: results of a mathematical model. Science Transl Med 1: 7ra16–7ra16.

43. Spencer SE, Besser TE, Cobbold RN, French NP (2015) ‘Super’ or just ‘above average’? Supershedders and the transmission of Escherichia coli O157: H7 among feedlot cattle. J R S Interface 12: p20150446.

44. Slater N, Mitchell RM, Whitlock RH, Fyock T, Pradhan AK, et al. (2016) Impact of the shedding level on transmission of persistent infections in Mycobacterium avium subspecies paratuberculosis (MAP). Vet Res 47: 1–12.

45. Kao RR, Gravenor MB, Charleston B, Hope JC, Martin M, et al. (20) Mycobacterium bovis shedding patterns from experimentally infected calves and the effect of concurrent infection with bovine viral diarrhoea virus. J R S Interface 4: 545–551.

46. Harvill ET, Cotter PA, Yuk MH, Miller JF (199) Probing the function of Bordetella bronchiseptica adenylate cyclase toxin by manipulating host immunity. Infect Immun 67: 1493–1500.

47. Levins R, Mohtashemi M (2001) Transient dynamics and early diagnostics in infectious disease. Journal of Mathematical Biology 43: 446–470.

48. Pugliese A, Gandolfi A (2008) A simple model of pathogen – immune dynamics including specific and non-specific immunity. Math Biosci 214: 73–80.

49. Fenton A, Perkins SE (2010) Applying predator-prey theory to modelling immune-mediated, within-host interspecific parasite interactions. Parasitology 137: 1027–1038.

50. Diniz LM, Magalhaes EF, Pereira FE, Dietze R, Ribeiro-Rodrigues R (2010) Presence of intestinal helminths decreases T helper type 1 responses in tuberculoid leprosy patients and may increase the risk of multi-bacillary leprosy. Clin Exp Immunol 161: 142–150.

